# Studying the role of climate and sex in flowering dynamics of black poplar across time and space

**DOI:** 10.64898/2026.10.05.756621

**Authors:** Cazenave Xabi, Villar Marc, Desforges Julien, Jorge Véronique, Duruflé Harold

## Abstract

Global change has a major impact on rivers and their associated terrestrial compartment, the riparian forest. The black poplar (*Populus nigra* L.), the dominant species of the European riparian forest, is considered an indicator of the biodiversity dynamics of these ecosystems, but is threatened with extinction in several parts of its range. The species is dioecious and shows wide variability in many traits, enabling it to adapt to a broad range of environmental conditions. A detailed characterization of this natural variability is therefore a key issue both for the conservation of natural populations and for biological engineering aimed at restoring riverbanks. Phenology is both an indicator of climate change and an adaptation lever for populations, particularly in perennial species such as trees. In this study, we characterized the spatial and temporal variability in flowering phenology by studying 13 natural black poplar populations along the Loire River, France. Flowering date and duration showed substantial variability both within and between populations, and males generally flowered earlier than females. Despite this variability, the ranking of individual trees’ flowering dates was highly consistent across years. Most male-female pairs shared several days of overlapping flowering each year, and this overlap was moderately stable across years. Flowering dates advanced by about five to six days per degree Celsius during early spring, with no detectable difference in sensitivity between sexes. This work contributes to the characterization of flowering variability in an economically important tree species and may inform both the conservation of genetic resources and breeding.

**Highlights:**

- Flowering phenology varied substantially within and among black poplar populations.
- Male black poplars generally flowered earlier than females.
- Flowering advanced by 5–6 days per °C increase in early spring temperature.
- Phenological diversity informs black poplar conservation and breeding.

## 1. Introduction

Phenology is the timing of recurring biological events that are influenced by physical environmental signals (Lieth, 1974) and includes several phenophases, or biological processes of development (*e.g.,* flowering time). Phenophases are sensitive indicators of the environment because they display strong responses to gradual changes in abiotic conditions, defining growth seasons in temperate forests. Phenology is also a pivotal process in tree adaptation because, as sessile organisms, trees are directly affected by their environment and rely on signals and regulations to synchronize biological events with changing conditions. Phenology plays a crucial role as one of the major adaptive traits in nature and acts as a bio-indicator of climate change. Phenological changes are related to phenotypic plasticity, reflecting how trees respond to environmental shifts. In the past decades, global climatic changes have disrupted phenological processes and produced a great acceleration of plant phenological shifts, which varies in magnitude between tree species (Fang et al., 2022; Geng et al., 2022; Menzel et al., 2020; Pausas, 2025; Piao et al., 2019; Vitasse et al., 2022). However, less is known about how interannual environmental variation affects the phenology of individual trees, especially flowering in natural populations.

The recent high tree mortality following extreme climate events, which occurs before maturity, raises concerns about the future of forests. The lack of knowledge makes it impossible to predict whether these observations represent a trend or to determine critical thresholds (Hartmann et al., 2022). Many studies in experimental conditions or in the field have found clinal patterns of phenology that are usually correlated with some environmental variables such as temperature, precipitation, latitude or altitude (Caicedo et al., 2004; Savolainen et al., 2007). For example, the northern limit of plant species ranges in the Northern Hemisphere appears to be driven by the inability to undergo full fruit maturation and the southern limit of plant ranges of the Southern Hemisphere seems to be driven by the inability to flower due to a lack of chilling temperatures that are necessary to break bud dormancy (Chuine, 2010). Moreover, long-term standardized monitoring networks such as the Swiss Commission for Phenology and Seasonality (KPS, https://kps.scnat.ch/en), the French TEMPO network (https://tempo.pheno.fr/eng), the USA National Phenology Network (USA-NPN, https://www.usanpn.org) and the Pan European Phenology PEP725 (Templ et al., 2026) underscore the importance of methodological consistency and replication in natural settings to ensure the comparability and robustness of climate-phenology inferences.

Intraspecific variability in phenological traits arises from both genetic differentiation among populations and phenotypic plasticity within populations. Common-garden studies in several tree species, including *Populus*, have shown that phenological traits are under genetic control and display clinal variation with latitude of tree origin (McKown et al., 2018). Interannual climatic variation also drives substantial plasticity in phenology in trees, with a recent large-scale analysis suggesting that plasticity, rather than local genetic adaptation, is the primary source of temperature-mediated variation in flowering time (Ramirez-Parada et al., 2024). This pattern may be even more pronounced in long-lived, wind-pollinated tree species (Liu et al., 2025). This predominance of plasticity may be particularly valuable given the inherently slow pace of genetic adaptation in trees, whose long generation times limit the rate at which populations can adapt to rapidly changing climatic conditions (Feng and Du, 2022; Jump and Peñuelas, 2005; Zhou et al., 2022). Phenotypic plasticity may therefore act as a critical buffering mechanism, allowing trees to track short-term climatic variation while genetic adaptation, if it occurs, unfolds over a much longer timescale (Nicotra et al., 2010). Monitoring phenological processes in natural tree populations is therefore essential to assess whether they can keep pace with climate change through plasticity alone, or whether active conservation intervention will be needed (Aitken et al., 2008).

However, the monitoring of the phenology of flowering and its genetic variation remains challenging in woody species, particularly due to the size of the experiments required, and the time to flowering age (*e.g.,* 10 years for poplar, 50 years for oaks). Beyond these logistical constraints, response to environmental cues is also harder to predict in dioecious species, in which it has been shown that the response could differ between males and females (Buonaiuto, 2024; Xie et al., 2023). In this regard, assessing the impact of climate change on both sexes is critical because a mismatch in flowering phenology between males and females could reduce fertilization success and endanger dioecious species (Hultine et al., 2016), even though there seem to exist stabilizing mechanisms in some dioecious species that constrain phenological shifts (Wu et al., 2025).

Among such species, poplar is a widely used model for plant biology, genomics, evolutionary and ecological genetics studies and has an economic and ecological interest (Douglas, 2017; Jansson and Douglas, 2007). Black poplar (*Populus nigra* L.), living on the banks of the river in the riparian zone of Europe, northwestern Africa, and Western Asia, has become a model species for *ex situ* and *in situ* conservation genetics (Lefèvre et al., 2001). Black poplar is clearly adapted to the riparian environment and its life cycle is affected at all stages by the hydrological conditions on which its regeneration depends (Corenblit et al., 2014).

Previous studies using microsatellite markers have shown high genetic diversity within populations and low genetic differentiation between river basins, suggesting high levels of gene flow in western parts of the distribution (Dewoody et al., 2015; Smulders et al., 2008). Resources for genetic studies of *P. nigra* have since been obtained, including SNP resources and the development of an Infinium 12k array providing a valuable tool for population genetic studies (Faivre-Rampant et al., 2016). Using this tool, the first genome-wide genotyping study was conducted on a large collection of native European black poplars (Allwright et al., 2016). Analyses of 838 native individuals from 12 river basins in Western Europe, based on a subset of 600 SNPs, identified seven ancestral genetic groups. Another study of seven European tree species showed that black poplar is genetically structured at the European level and displays moderate genetic diversity compared to other tree species (Milesi et al., 2024).

Trees from the *Populus* genus are dioecious and therefore have distinct unisexual individuals. These species do not have sex-specific heteromorphic chromosomes (Tuskan et al., 2012) but only a single sex-determining gene (Müller et al., 2020) which undergoes epigenetic regulation (Bräutigam et al., 2017). Some studies show sexual dimorphisms in poplar before and after sexual maturity, such as growth for *P. yunnanensis* (Lei et al., 2017; Zhang et al., 2019), photosynthesis and water status for *P. cathayana* (Liu et al., 2020), or more broadly the response to environmental stresses among the *Populus* genus (Melnikova et al., 2017). In contrast, no evidence of sexual dimorphism has been found in flowering phenology or differential resource investment strategies between males and females in either sexually immature or mature *P. tremula* trees (Robinson et al., 2014). Sexual homomorphism has also been observed for several traits in North American populations of *P. trichocarpa* and *P. balsamifera*, suggesting that sex could be functionally neutral concerning non-reproductive traits that might affect plant survival (McKown et al., 2017). Even though sexual dimorphism seems trait and species-specific in the *Populus* genus, males and females in a given species can be expected to differ in phenology because their resource allocation strategies differ. Indeed, males invest their resources in pollen, before anthesis, while females allocate more resources during seed production after anthesis (Barrett and Hough, 2013). A desynchronization of flowering time of the two sexes in response to climate change could affect the reproductive success and thus the diversity of the species (Buonaiuto, 2024). In this context, (Xie et al., 2023) estimated different phenological sensitivities between males and females for eight poplar species distributed across North America, but black poplar was not one of the studied species.

The present study thus aimed to characterize the interannual and interpopulation variability of flowering phenology for both sexes in black poplar. Flowering phenology was assessed in individuals from 13 natural populations distributed along the Loire River, spanning a latitudinal and climatic gradient across France. One of these populations was monitored over a period of 22 years. The long-term monitoring of this population allowed us to assess the influence of climatic drivers on flowering phenology, as well as to investigate potential sexual dimorphism in phenological timing and response to climate change.

## 2. Material and methods

### 2.1 Sites and tree material

Thirteen natural populations of black poplar (*Populus nigra* L.) were studied along the French Loire River (Fig.1A). The number of trees monitored in each population varied from 30 to 56 (Table 1) totalling 500 trees. Flowering phenology could be monitored for 482 of these trees. The proportion of females among the 500 trees was 0.42 and varied from 0.3 to 0.63 in the studied populations.

**Figure 1.**
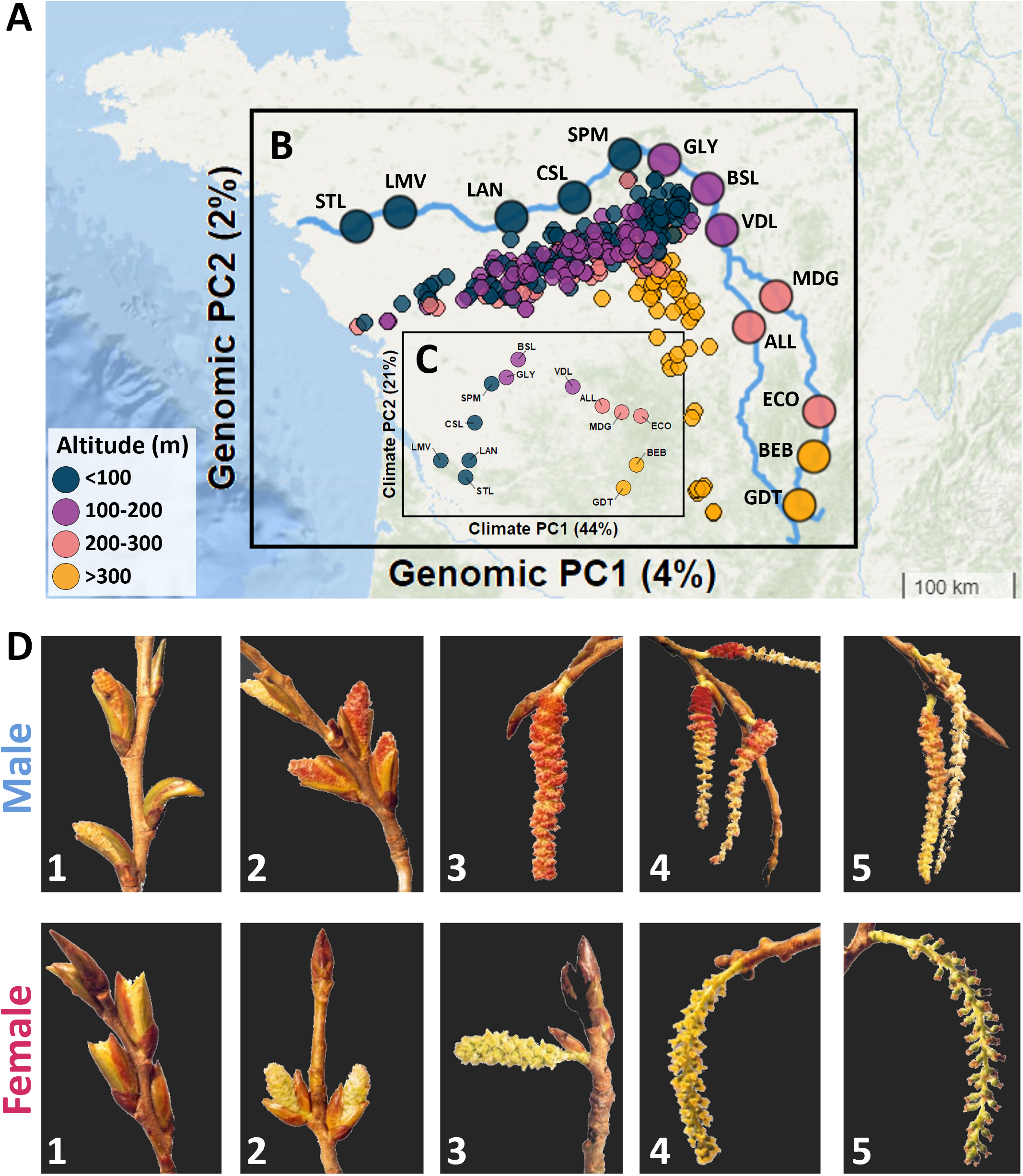
**A**. Overlay of a map of the French Loire River with **B.** the results of a principal component analysis (PCA) using 2,349 SNPs from 445 individuals from 13 natural populations coloured according to their altitude. Each point represents one of the 500 individuals plotted according to their PC1 and PC2 values. **C.** the results of a PCA using climatic variables from the 13 specific locations. **D**. Phenological stages used to monitor poplar flowering. Phenological scale was defined as follows: 0 - Dormant bud; 1 - Bud sprouting, with tips of early flowers emerging out of the scales; 2 - Bud sprouting, with half of the flowers of the catkin emerging out of the scales; 3 - Bud completely open, with flowers still clustered together. Red stamen / yellow stigmas flowers are easily visible. Stigmas flowers are turgescent; 4 - Bud open, full lengthening of the axis of the catkin. Male flowers shed pollen / stigmas of the female flowers are turgescent and receptive; 5 - Catkins have shed all the pollen / stigmas become brown and are not receptive any more.

**Table 1:** Characteristics of the natural French Loire River populations follow for this study.

| Acronym | Location | n | Sex ratio | Latitude | Longitude | Altitude (m) |
| --- | --- | --- | --- | --- | --- | --- |
| ALL | Val d'Allier | 45 | 0.38 | 46.4000 | 3.3167 | 227 |
| BEB | Bas en Basset | 35 | 0.63 | 45.3067 | 4.1100 | 470 |
| BSL | Bonny sur Loire | 41 | 0.39 | 47.5667 | 2.8167 | 133 |
| CSL | Chaumont sur Loire | 33 | 0.30 | 47.4808 | 1.1864 | 85 |
| ECO | Ecopole du Forez | 36 | 0.36 | 45.6978 | 4.1772 | 334 |
| GDT | Goudet | 30 | 0.33 | 44.8902 | 3.9222 | 826 |
| GLY | Guilly | 39 | 0.39 | 47.8000 | 2.2833 | 108 |
| LAN | Langeais | 35 | 0.34 | 47.3267 | 0.4069 | 61 |
| LMV | Le Mesnil en Vallée | 33 | 0.46 | 47.3667 | -0.9342 | 39 |
| MDG | Méandre des Germaines | 35 | 0.34 | 46.6686 | 3.6567 | 202 |
| SPM | Saint-Pryvé-Saint-Mesmin | 51 | 0.56 | 47.8500 | 1.8000 | 95 |
| STL | Sainte-Luce | 31 | 0.52 | 47.2534 | -1.4828 | 15 |
| VDL | Val de Loire | 56 | 0.41 | 47.2167 | 2.9833 | 153 |

n: Number of individuals per population

### 2.2 Monitoring of flowering

A population of 51 trees located in Saint-Pryvé-Saint-Mesmin (SPM, 47°51’39.6“N 1°46’26.4”E) was monitored annually between 2003 and 2024. Years with an insufficient number of scoring dates to reliably estimate flowering phenology were excluded, retaining seven years of data: 2003, 2004, 2006, 2013, 2022, 2023, and 2024. In addition, 12 other populations located along the Loire River were monitored in 2006 using the same scoring method as the one used in SPM. In both cases, each tree was monitored at least once a week (and up to three times a week) and was scored between 0 and 5 following the scale presented in Figure 1D. Briefly, the phenological scale was defined as follows: 0 - Dormant bud; 1 - Bud sprouting, with tips of early flowers emerging out of the scales; 2 - Bud sprouting, with half of the flowers of the catkin emerging out of the scales; 3 - Bud completely open, with flowers still clustered together. Red stamens / yellow stigmas are easily visible. Stigmas of the female flowers are turgescent; 4 - Bud open, full lengthening of the axis of the catkin. Male flowers shed pollen / stigmas of the female flowers are turgescent and receptive; 5 - Catkins have shed all the pollen / stigmas become brown and are not receptive any more.

### 2.3 Genotyping and genomic data analysis

Out of the 500 monitored trees, 314 were genotyped with the 12k Infinium array described in(Faivre-Rampant et al., 2016) and 131 trees were genotyped with a multi-species Axiom SNP array, including 13,408 SNPs for poplar, developed during the B4EST European project (Guilbaud et al., 2026). Missing data imputation was carried out using the BEAGLE 5.5 software (Browning et al., 2018) with 100 burn-in iterations followed by 1000 iterations. In total, 2,349 SNPs were used both to confirm the absence of clonal duplicates among the monitored trees and to characterize the genetic structure of the studied populations through principal component analysis (PCA), performed on allele dosage data.

### 2.4 Climatic data

The Climate Downscaling Tool geo-web service (Marchi et al., 2020) was used to extract monthly and annual weather variables from the 13 populations between 2002 and 2024 with a resolution of *ca.* 1 km^2^ per grid cell. Daily minimum and maximum temperatures were also downloaded from the ERA-5 database (Muñoz-Sabater et al., 2021) for the same GPS coordinates and time period. These variables were extracted and downscaled using the KrigR package v0.9.0 (Kusch and Davy, 2022). For subsequent analyses, a phenological season running from September of year n (corresponding to bud formation) to April of year n+1 (corresponding to the end of flowering) was defined and data from the May-August period were discarded. Additionally, we extracted the 19 bioclimatic parameters (bio1 to bio19) from the WorldClim v.2.1 database (Fick and Hijmans, 2017) for each of the 13 populations for the reference period of 2006, during which all populations were monitored, and used the values obtained at the population level to run a PCA and characterize each population in terms of climate.

### 2.5 Flowering data analysis

Flowering phenology was modelled using a linear regression for each tree monitored, using the recorded phenological stages as outcome variable and days of the year as explanatory variable. Three phenological parameters were extracted from the linear regression output: beginning of flowering, which corresponds to the estimated score 1 date as a Day Of Year (DoY); end of flowering, which corresponds to the estimated score 5 DoY, and flowering duration, which is the difference between the end and beginning dates.

### 2.6 Synchrony between males and females

Common flowering period was computed for all the male/female pairs in the 13 populations and the 7 studied years in SPM population, where the common period represents the total number of days during which both the male and the female exhibited flowers. A heatmap of all the pairs was then computed using the Optimal Leaf Ordering (OLO) clustering algorithm (Bar-Joseph et al., 2001). To assess the year-to-year stability of the flowering period overlap within the SPM population, we fitted a linear mixed model with tree pair as a random intercept and year as a fixed effect, and computed the adjusted intraclass correlation coefficient (ICC) as the proportion of variance explained by the between-pair differences after accounting for year effects.

The year-to-year consistency of pairwise common flowering duration was also assessed with Spearman correlations between years. In addition, for each pair of years, we calculated the proportion of pairs with less than 3 days of common flowering (or in the upper quartile of common flowering duration) in the first year that remained in the same category in the second year, and compared it with the proportion of all pairs in that category in the second year.

### 2.7 Correlation between climatic variables and phenological parameters

In order to identify the climatic variables that are the most related to the phenological parameters, we computed the correlation between each climate variable and the studied phenological parameters. To identify the period of the year during which mean daily temperature most strongly affects flowering phenology, a separate Partial Least Squares (PLS) regression was fitted for each sex with the chillR R package (Luedeling et al., 2013a), following the approach of (Luedeling and Gassner, 2012). Each model related the annual mean date of the beginning or end of flowering, averaged across all monitored trees of that sex, to daily mean temperatures from September of the previous year to April, over the seven monitored years. The threshold for the variable importance in the projection (VIP) is commonly set to 0.8 (Guo et al., 2013), so we adopted this threshold for our analyses.

### 2.8 Temperature sensitivity

To compare the response of males and females to temperature, we used the mean temperature over the period identified by these PLS models as most strongly associated with flowering (see Results). For each year, the temperature anomaly was the difference between this mean and its average over the seven monitored years. A linear mixed model was fitted separately for the beginning, end and duration of flowering, with sex, temperature anomaly and their interaction as fixed effects and tree and year as random intercepts. Fixed effects were tested with F-tests using the Kenward-Roger approximation from the *mixed()* function from afex R package, and the variance explained was estimated with marginal and conditional R² (Nakagawa and Schielzeth, 2013).

Following (Xie et al., 2023), phenological sensitivity was defined as the change in flowering date (in days) per °C of temperature anomaly, i.e. the slope of the temperature anomaly in the model. The sensitivity of each sex and the difference between sexes were calculated as the estimated slopes of the temperature anomaly for females and males with the *emtrends()* function in the emmeans R package (Lenth et al., 2017).

### 2.9 Statistical software and data visualization

All statistical and genetic analyses were run using `R` v4.5.1 (R Core Team, 2021) under the rstudio environment (RStudio Team, 2020). The map presented in Figure 1 was created using the `leaflet` R package v2.2.2 (Cheng et al., 2025) and PCA was performed with the R package `mixOmics` v6.28.0 (Rohart et al., 2017). The linear mixed models were run using the `lmer` function of R package `lme4` v.1.1.37 (Bates et al., 2015).

## 3. Results

### 3.1 Genetic and climatic structure across the Loire basin

To assess the natural genetic diversity of black poplar along the Loire River, we sampled 13 populations distributed along the entire length of the river, from the estuary (STL population) to the source (GDT population) (Figure 1A). A random selection of at least thirty genotypes per population was performed (Table 1).

A visualization of the genomic variation of the 445 selected individuals using principal component analysis (PCA) on SNP data revealed no differentiation between the Loire River populations, except for the populations at higher altitudes (Figure 1B). Indeed, populations located at altitudes below 300 m appeared to be poorly structured and distributed along the PC1, while populations at higher altitudes, GDT and BEB, appeared to be more distinct along the PC2. A coefficient of determination (R²) of 0.55 was observed between altitude and the genetic PC2 (Figure S1A).

The multivariate analysis of 19 bioclimatic variables from the WorldClim portal using the Climate Downscaling Tool geo-web service was carried out via PCA for the year 2006 (Figure 1C). Using this approach, a nonlinear environmental gradient was observed between the PC1 and PC2 axes, as a reflection of the geographical location of populations. Additionally, a coefficient of determination of 0.189 was observed between the genetic PC1 and the PC2 obtained from climatic data (Figure S1B).

### 3.2 Timing of flowering in males and females

A linear regression of the phenological stage over day of year (DoY) was performed for each of the 482 monitored individuals. This approach allows us to extrapolate unobserved stages and also to extract three phenological parameters: the beginning, the end and the duration of flowering.

In 2006, the beginning of flowering for the 11 lowland populations (below 300 m altitude) happened between the 70th and 105th day of the year, while flowering started later for the BEB and GDT populations, i.e. between the 90th and 120th day of the year. There was no statistical difference between males and females for the beginning of flowering for 7 out of the 13 studied populations (Figure 2A). When the difference between sexes was significant (i.e. for the STL, LMV, LAN, VDL, MDG and BEB populations), females always flowered later than males. Across all the studied populations, male individuals exhibited on average 3.0 +/- 0.4 days earlier flowering than females (P < 0.001).

**Figure 2.**
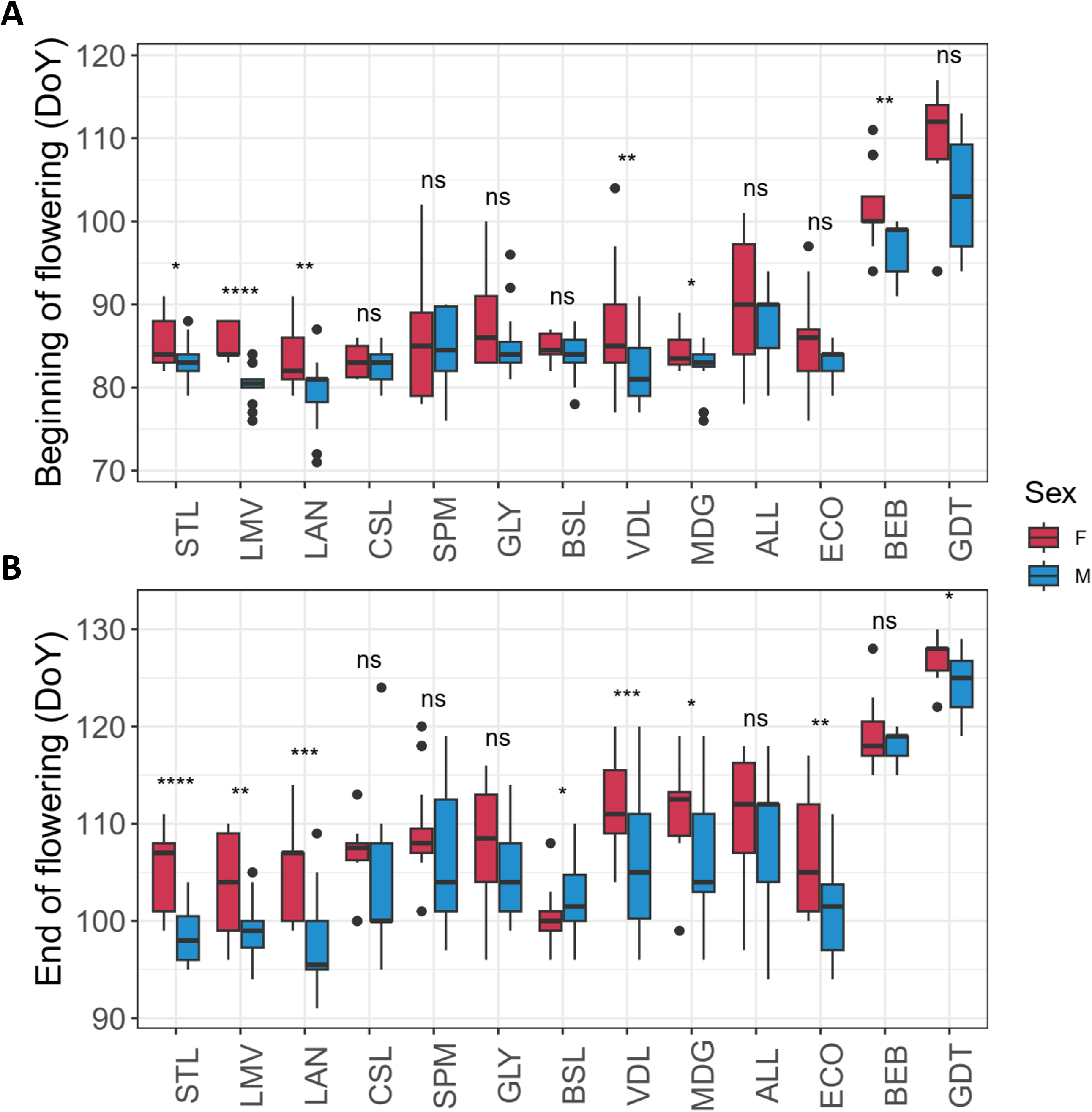
Boxplots of **A**. the beginning and **B**. the end of flowering between males and females from the 13 populations. (DoY: Day of year)

The end of flowering showed a greater variability in response (Figure 2B), but the same trend was observed, as females tended to end flowering later than males (on average 4.8 +/- 1.2 days later). This difference was significant in most populations, except for CSL, SPM, GLY and ALL. As a consequence, the duration of the female flowering period was generally longer than for males (mean duration in males: 15.8 days, mean duration in females: 17.1 days). Interestingly, we observed the opposite for the high-altitude populations (mean male and female flowering duration was 16.1 and 13.3 days respectively in BEB, 17.7 and 14.7 respectively in GDT).

As the SPM population was monitored over seven years (between 2003 and 2024), we also focused on this specific population to assess if the same trends could be observed over time. Considering all the studied years in SPM, male individuals flowered on average 2.5 +/- 0.3 days earlier than females. However, this pattern was not observed for every studied year: in 2003, 2006, 2013 and 2022, male and female beginning of flowering was not statistically different (Figure 3A). In contrast, regarding observations made over the years for the SPM population, we observed little variation between sexes for the end of flowering, except in 2006, for which male genotypes completed flowering earlier than female genotypes (Figure 3B).

**Figure 3.**
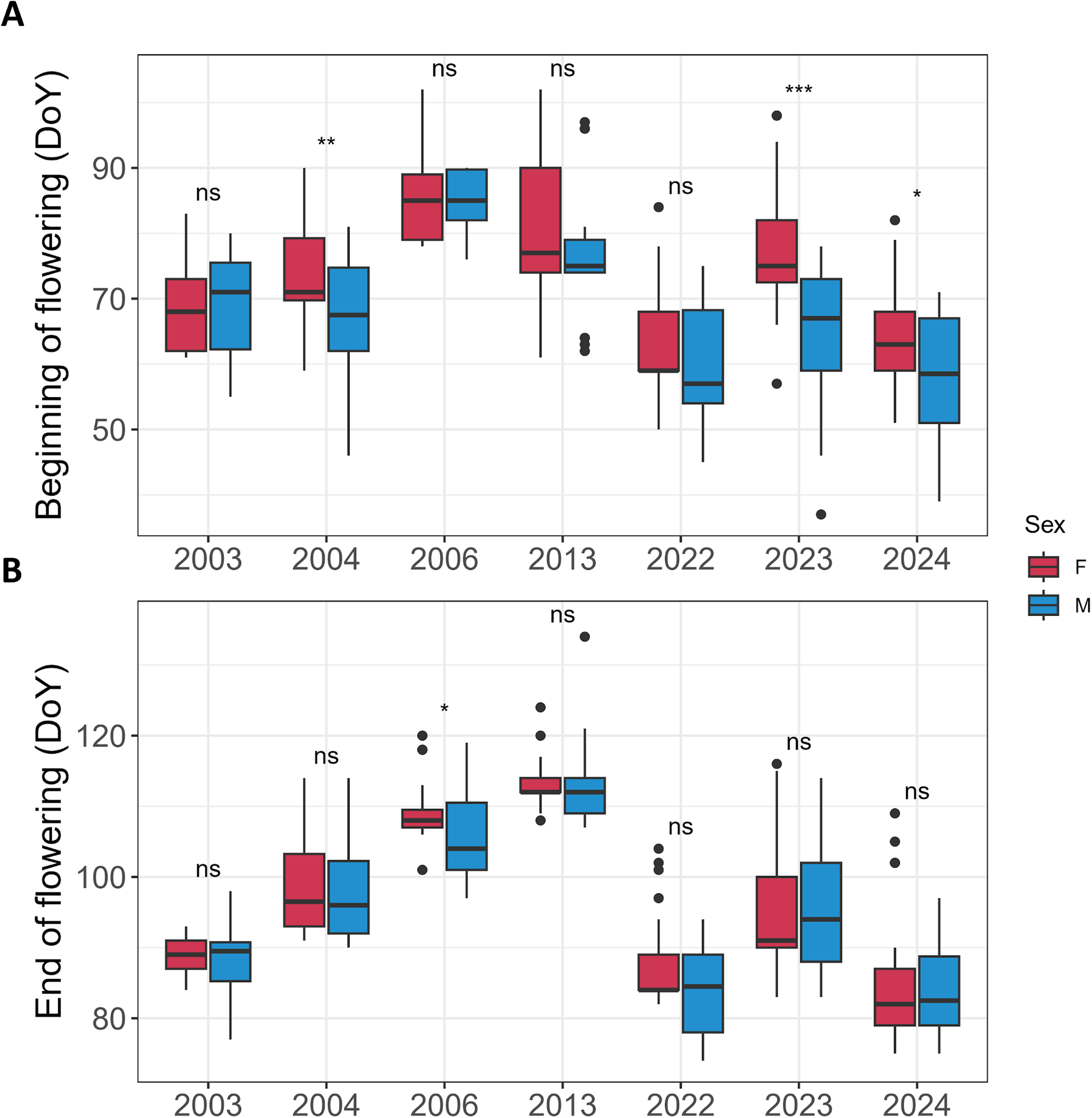
Boxplots of **A**. the beginning and **B**. the end of flowering between males and females from the SPM population for the monitored years. (DoY: Day of year)

### 3.3 Correlation in ranking of flowering period over years

To assess the variability of flowering dates across years, we focused on the SPM population, which is the only population out of the 13 studied ones to have been monitored during several years. Even though a substantial variability could be observed for the beginning of the flowering period for both males and females in SPM, we observed a strong correlation when considering the ranking of individual trees based on the flowering dates (Figure 4, Figure S2). The average Kendall correlation for the beginning of flowering was 0.78 and 0.83 for the end of flowering. This correlation was very high in both sexes (Kendall tau = 0.76 for beginning of flowering in males and 0.77 in females; Kendall tau = 0.82 for end of flowering in males and 0.81 in females, Figures S3 & S4). The correlation for flowering duration was nevertheless close to 0 across years (Figure S5), although substantial variability could be observed between year pairs (min: -0.08; max: 0.58).

**Figure 4.**
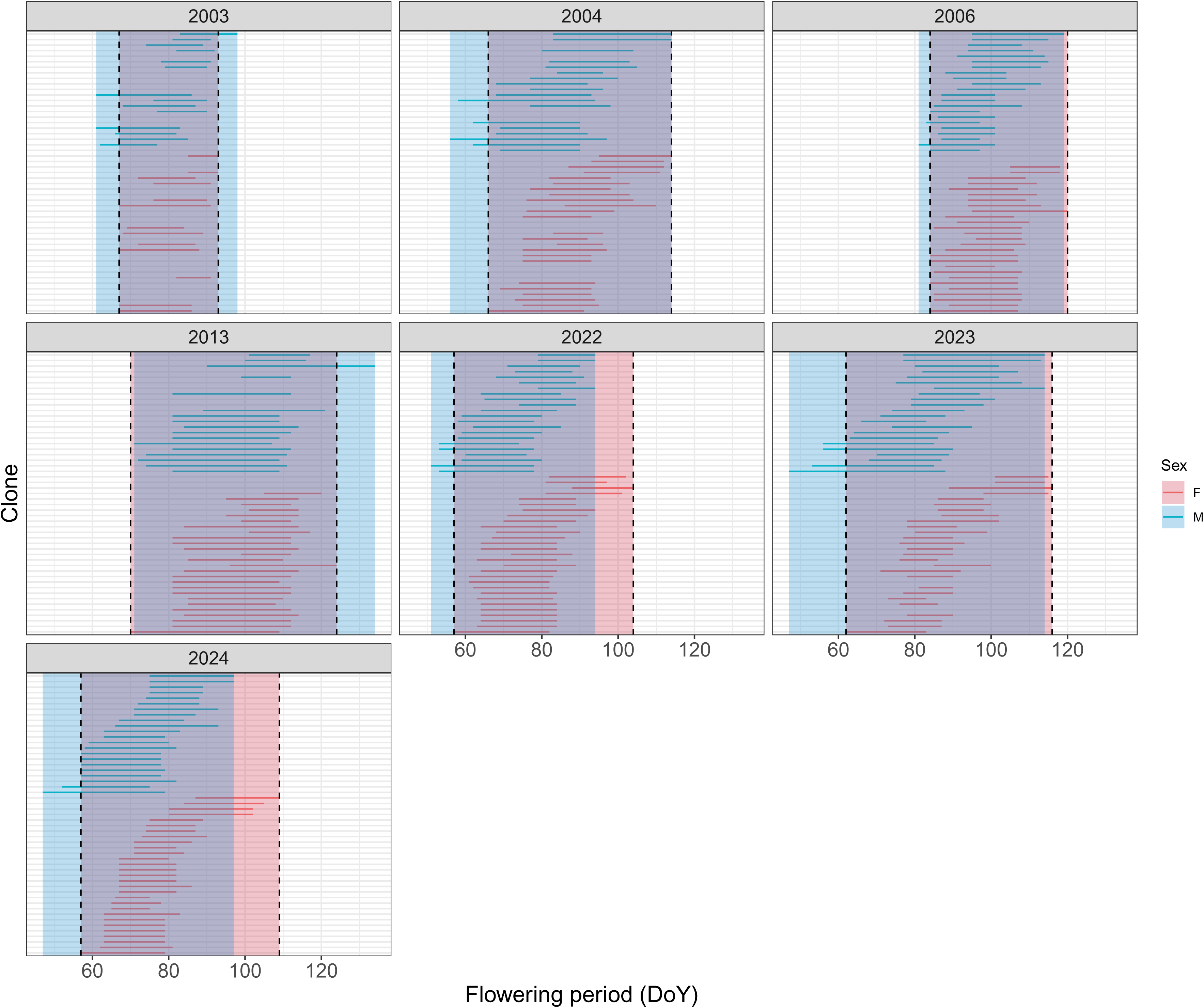
Flowering period by clone and across years in the SPM population. Each plain line represents a clone (females in red, males in blue) and the dotted lines correspond to the beginning and end of the common flowering period (i.e., the period for which at least one male and one female are flowering).

### 3.4 Flowering synchrony between sexes

To assess whether specific male-female pairs consistently exhibited similar durations of flowering overlap across years, we computed the adjusted intraclass correlation coefficient for pairwise common flowering duration for the SPM population, which indicated a moderate level of consistency (ICC = 0.53). Pairwise common flowering duration was also moderately correlated between years (Spearman ρ = 0.15–0.67, mean 0.47), without decline with the interval between years. Pairs with less than 3 days of common flowering in one year tended to have less than 3 days in other years as well. For example, among the 56 pairs with less than 3 days of common flowering in 2022, 79% also had less than 3 days in 2023, compared with 17% of all pairs in 2023. Across all combinations of years, a median of 63% of the pairs with less than 3 days of common flowering in one year also had less than 3 days in another year, compared with a median of 7% of all pairs. The same held, to a lesser extent, for the pairs with the longest common flowering periods: a median of 48% of the pairs in the upper quartile of common flowering duration in one year were also in the upper quartile in another year, compared with a median of 31% of all pairs. The maximum pairwise duration of common flowering varied among years, ranging from 19 days in 2003 to 37 days in 2013, with most years falling between 22 and 25 days (Figure S6). The large majority of male–female pairs had at least some degree of flowering overlap: across years, 83–100% of pairs shared at least 3 days of common flowering, and pairs with no overlap at all remained rare (0–12.5% depending on the year). The proportion of pairs with less than 3 days of common flowering appeared to increase over the most recent years monitored, from 8.8% in 2022 to 17.2% in 2023 and 10.5% in 2024, suggesting a possible recent trend toward greater desynchronization that would warrant confirmation with additional years of monitoring. Nearly all individuals had a common flowering period of at least 3 days with at least half of the opposite sex each year (100% of males and between 89 and 100% of females). When considering the 13 populations monitored in 2006, pairwise common flowering duration ranged from no overlap for a few pairs to 28 days (Figure S7). The proportion of pairs sharing at least 3 days of common flowering varied among populations, from 83% in ALL to 100% in BSL and CSL, and pairs with no overlap at all ranged from 0% to 13% of pairs depending on the population (Figure S7).

### 3.5 Impact of temperature on flowering phenology

The climatic variables which showed the highest correlation with phenological parameters were always spring temperatures, while precipitation or humidity variables showed moderate to low correlations to the phenological variables. We thus chose to run a PLS analysis based on mean daily temperatures to explore more precisely the period of the year that could affect phenological events in the SPM population. For both sexes, the direction of the effects remained stable over continuous periods. A first advancing period, spanning February to late March, was identified for the beginning of flowering (Figure 5), while a slightly later period, from mid-February to late March, was identified for the end of flowering. A second, shorter period of delaying effects was identified in early-to-mid October (about two to three weeks), of comparable magnitude to the advancing periods but of opposite direction.

**Figure 5.**
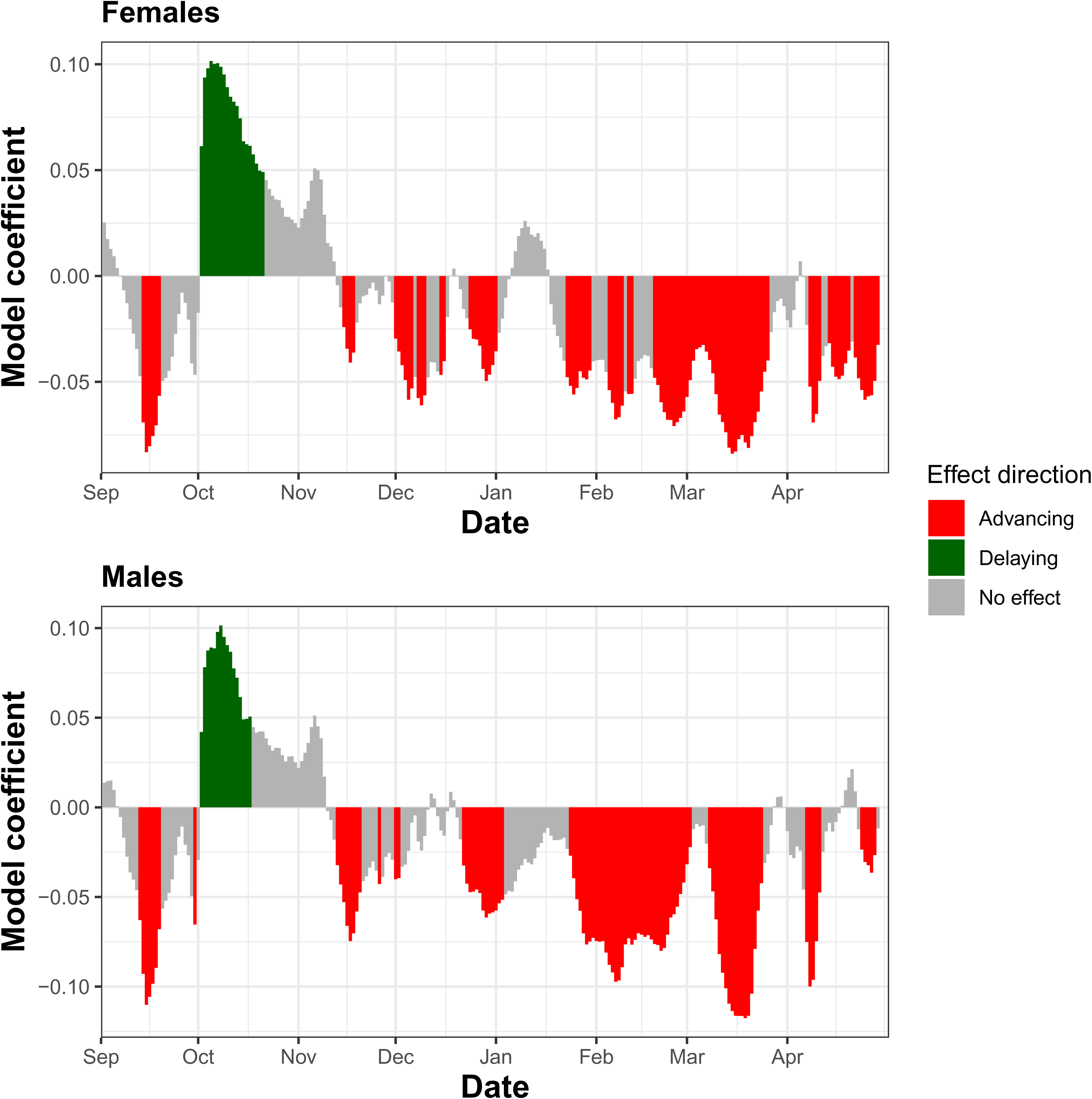
Effect of monthly mean temperatures on beginning of flowering in SPM for the females and the males. Bar plot in red represent days for which an increase in temperature leads to advancing of flowering and in green, days for which an increase in temperature leads to a delaying of flowering. The coefficients correspond to the estimated daily effects from the PLS model for individual trees.

Based on these results, we computed the average daily temperature for the period from February to March, common to both sexes and to the beginning and end of flowering, and used it as the climatic predictor in subsequent models. Temperature had a significant effect on the beginning and end of flowering, but not on its duration (Table 2). Sex also had a significant effect on the beginning and duration of flowering, but not on its end: females began flowering 4.7 days later than males (95% CI: 0.5 to 8.9) and had a 3.4-day shorter flowering duration (95% CI: −4.7 to −2.0). The phenological sensitivity of the beginning of flowering was −4.8 days/°C in females (95% CI: −7.3 to −2.3) and −5.2 days/°C in males (95% CI: −7.7 to −2.7), and that of the end of flowering was −6.1 days/°C (95% CI: −9.2 to −3.0) and −6.0 days/°C (95% CI: −9.0 to −2.9), respectively. Sensitivity did not differ between sexes: the interaction was not significant for any parameter, and the difference between females and males was small (beginning: 0.4 days/°C, 95% CI: −0.1 to 1.0; end: −0.1, −0.6 to 0.4; duration: −0.5, −1.2 to 0.2). Sex, temperature and their interaction explained 47% of the variance in the beginning of flowering, and 89% when tree and year were also included. For the end and duration of flowering, they explained 57% and 13% of the variance, respectively, and 93% and 40% when tree and year were also included.

**Table 2:** Effect of spring temperature anomaly (February–March), sex, and their interaction on the beginning, duration and end of flowering, with the resulting temperature sensitivity by sex and the variance explained by each model.

| Effect | Beginning | Duration | End |
| --- | --- | --- | --- |
| **Sex** | 5.01, p = 0.030 | 24.17, p < 0.001 | 0.66, p = 0.421 |
| **Temperature** | 26.36, p = 0.004 | 1.50, p = 0.275 | 25.42, p = 0.004 |
| **Sex × Temperature** | 2.24, p = 0.136 | 2.07, p = 0.152 | 0.18, p = 0.668 |
| **Sensitivity, females (days/°C)** | -4.8 [-7.3, -2.3] | -1.3 [-3.4, 0.8] | -6.1 [-9.2, -3.0] |
| **Sensitivity, males (days/°C)** | -5.2 [-7.7, -2.7] | -0.7 [-2.9, 1.4] | -6.0 [-9.0, -2.9] |
| **Marginal R<sup>2</sup>** | 0.47 | 0.13 | 0.57 |
| **Conditional R<sup>2</sup>** | 0.89 | 0.40 | 0.93 |

## 4. Discussion

### 4.1 Genetic structure and flowering phenology across the Loire basin

In the Loire River basin, the genetic structure of populations appeared weak, and no differentiation was observed except for high-altitude populations. This differentiation is likely due to local adaptation in colder environments resulting in reduced gene flow with the other populations. Flowering dates showed substantial variability within and between populations. The distribution of flowering dates within populations approximately followed a normal distribution centred on a period of fertility, suggesting stabilizing selection around a locally optimal reproductive period as has been observed in other perennial species (Gauzere et al., 2019). At the interpopulation level, flowering phenology broadly mirrored this pattern of weak structure: the low-altitude populations, genetically undifferentiated from one another, also flowered at similar dates, while the two high-altitude populations, genetically distinct, also flowered later — potentially reflecting genetic differentiation, colder conditions, or both. This is consistent with a growing number of recent genomic-environment association studies linking phenological divergence to adaptive variation along climatic clines (Lazic et al., 2024).

These trends are consistent with previous work documenting spatial genetic structuring and adaptive differentiation at regional scales and align with recent large-scale analyses integrating phenology and population structure at comparable spatial resolutions (Żukowska and Lewandowski, 2025). Collectively, these observations support a scenario in which weak overall structuring across the Loire is punctuated by altitude-related differentiation, while intrapopulation phenological variation and interpopulation phenological divergence are jointly shaped by climatic gradients (temporal or spatial) and, potentially, genetic differentiation.

### 4.2 Impact of Climatic Variables on Flowering Phenology

Identifying the climatic drivers of flowering phenology, and more specifically the periods of the year during which they exert the strongest influence, is essential to anticipate how tree populations may respond to ongoing climate change and to inform conservation strategies. However, disentangling the effect of specific climatic windows from the broader seasonal temperature signal remains statistically challenging. Partial least squares (PLS) regression allows the identification of climatic factors influencing plant phenology when the explanatory variables are highly collinear and have a fine temporal resolution. This situation is typical of temperature time series, where daily or weekly measurements are highly autocorrelated and often more numerous than phenological observations. PLS regression has been shown to reliably identify biologically relevant temperature windows influencing phenological events (Luedeling et al., 2013b; Luedeling and Gassner, 2012; Ranjitkar, 2013), which makes it well suited in principle to the type of data analysed here. In our analysis, temperature variables, particularly those characterizing early spring, emerged as the main predictors of flowering time, while precipitation variables did not show significant explanatory power.

The predominance of temperature effects is consistent with the well-established role of heat accumulation in regulating budburst and flowering in temperate trees (Basler and Körner, 2012; Cleland et al., 2007). Recent large-scale syntheses confirm that late winter and early spring temperatures exert control over phenological transitions, especially in the context of current climate warming (Pandey et al., 2026). In poplars, flowering and bud development are tightly regulated by chilling requirements followed by heat accumulation, with a high sensitivity to temperature anomalies in late winter (Ibáñez et al., 2010; Rohde et al., 2011). Furthermore, the flowering period in poplars is a heritable trait, with stable inter-individual rankings from year to year, confirming the existence of a significant genetic component underlying phenological variation (Mckown et al., 2014).

The absence of detectable water effects could reflect the ecological specificity of the studied system. Poplars are characteristic species of riparian environments, generally found in areas where soil moisture is relatively stable and high. In these habitats, water limitation is less likely to act as a phenological factor than temperature (Alimpić et al., 2022). Indeed, although water stress can influence growth and phenology in trees, most studies on spring phenology identify temperature as the primary determining factor, with water variables playing a secondary or indirect role, except in cases of severe water deficit (Kalcsits et al., 2009; Soolanayakanahally et al., 2013). This capacity for ecological regulation in response to water variability could therefore explain why only thermal predictors were identified in our models.

The longitudinal monitoring of a natural population is a major strength of this study. By repeatedly assessing the same population over several years, we effectively minimize confounding effects related to interpopulation genetic differentiation, allowing us to more clearly attribute interannual variation to climatic variables rather than to local adaptation possibly present within populations. Long-term phenological monitoring within a single population has been shown to be a powerful tool for distinguishing plastic responses from spatial genetic differentiation (Anderson et al., 2012). However, our dataset also has limitations. Notably, gaps in temporal coverage and a relatively small number of consecutive years restrict our ability to test for delayed or cumulative climatic effects from year to year, approaches that have recently revealed complex, persistent, and heritable effects in perennial species (Blonder et al., 2023). Extending this long-term monitoring effort in the coming years would therefore be particularly valuable to test for such delayed or cumulative stress effects, and more broadly to strengthen our ability to detect and anticipate phenological responses to ongoing climate change.

### 4.3 Sex-specific flowering timing and its response to climate

Flowering phenology in *Populus* is not only heritable, but also potentially structured by sex-specific dynamics. Preferential synchrony between male and female individuals within populations has been observed in dioecious tree species and may influence pollen circulation and local mating patterns of flowering phenotypes (Bogdziewicz et al., 2020; Xie et al., 2023).

Phenological differences between males and females in flowering time are also well documented in dioecious plant species including for the genus *Populus* (Vanden Broeck et al., 2003), where males often flower earlier than females (a phenomenon known as protandry), which promotes long distance pollination (Ou et al., 2025). In this context, a large-scale study demonstrated earlier flowering in males, as well as a sex-specific sensitivity to climatic factors (Xie et al., 2023).

In line with these observations, males in SPM began flowering earlier than females on average (by 4.7 days), consistent with the protandry reported in other *Populus* species. However, we found no evidence that males and females differed in their sensitivity to spring temperature, suggesting that, while the two sexes flower at different times, their phenological response to interannual climate variation is similar in this population. The phenological sensitivity to spring temperature observed in SPM (4.8–6.1 days °C ¹ depending on sex and phenological parameter) was higher than the sensitivities reported by Xie et al. (2023) for eight North American *Populus* species (around 2 days °C ¹). This discrepancy likely reflects both methodological and study design differences: our temperature window was selected using the same seven years subsequently used to estimate sensitivity, and herbarium-based estimates such as those of Xie et al. are prone to measurement error that would tend to attenuate slope estimates. In addition, our estimates rely on only seven years of data from a single population, whereas Xie et al. drew on multiple populations and decades of records, which likely yields more robust and generalizable sensitivity estimates. Conclusions on sex-specific climate sensitivity in SPM should therefore be treated with some caution given these constraints.

Beyond average phenological differences, we observed that specific male-female pairs tended to consistently show either high or low flowering overlap across years: pairs with less than three days of common flowering in a given year were considerably more likely to remain in this category in another year than pairs in general and a similar (albeit weaker) pattern was found for pairs with the longest overlap. This pairwise consistency (ICC = 0.53) is at least partly explained by the fact that each tree’s own flowering phenology remains relatively stable across years. Such stability could influence effective gene flow, effective population size, and genetic diversity profiles. The ecological and evolutionary implications of sex-specific phenological structuring are potentially considerable. Temporal segregation between males and females can modify reproductive match, while also modulating patterns of sexual selection and adaptive potential (Slavov et al., 2025).

Even subtle differences in the duration or timing of flowering could alter effective sexual overlap, thereby influencing mating networks and genetic diversity. Comparison between the SPM population and all populations along the French Loire River suggests that sex-specific flowering patterns in *Populus* may be more context-dependent than previously thought. Determining whether this similarity in climate sensitivity between sexes observed in SPM (and thus the likelihood that the timing gap between males and females remains stable under further warming) can be generalized to other populations will require larger multi-year, multi-population datasets that integrate phenological analysis, spatial structure analysis, and genetic relatedness analysis. Such efforts are essential to assess the ecological consequences of a possible temporal separation between males and females on the dynamics of pollen flow and the long-term resilience of populations to climate change.

## 5. Conclusions and perspectives

To our knowledge, this study is the largest one reported in terms of number of populations, trees and years of observations on flowering phenology in *Populus nigra*. Our work highlights the predominant role of February-March temperatures in the flowering phenology of natural riparian poplar populations. The relative phenological stability observed from year to year, combined with consistent differences in flowering timing between males and females, suggests that reproductive overlap between specific individuals tends to remain stable across years. If phenological shifts alter male-female synchronization or effective reproductive periods, consequences for pollen flow, effective population size, and genetic diversity could emerge. Previous work on dioecious trees has demonstrated that climate-induced changes in sex ratio or reproductive resource allocation can affect population stability and genetic structure (Petry et al., 2016).

However, the limited number of years of monitoring and individuals studied restricts the robustness of the conclusions regarding interannual climate sensitivity, delayed effects, and sex-specific responses. Therefore, extending the observation period and increasing the sample size will be essential to detect potential nonlinear or cumulative climate effects, which are recognized as crucial components of the phenological dynamics of perennial plants (Grossman, 2023). Future work should also refine phenological monitoring methodologies. Automated or semi-automated sensor-based approaches, including high-frequency imaging and environmental proximity sensors, would improve temporal resolution and reduce observational biases, particularly in systems where phenological transitions are nonlinear and sensitive to short-term temperature variations.

Integrating long-term phenological monitoring, spatial genetic analyses, and lineage reconstruction would thus allow for a better understanding of whether observed phenological stability, or potential future divergence, translates into measurable variations in genetic diversity. Such an approach to monitoring a single natural tree population provides a valuable model for examining the combined ecological and evolutionary consequences of climate-induced phenological changes in riparian forest trees.

## Supporting information

Supplementary figure S1 - Correlation between the altitudes, genetic values and climatic PC2 for each individual

## Acknowledgements

The authors would like to warmly thank the managers of the natural sites, who gave access to the trees and who participated in the collection of the 2006 data: Christophe Tomati in GDT, Philippe Cochet in BEB, Diane Corbin in ECO, Daniel Mayereau in MDG, Nicolas Pointecouteau in VDL, Michel Chantereau in BSL, GLY and CSL, Michel Chantereau and Damien Hémeray in SPM, Yves Gabory and Ambroise Bécot in LMV, Xavier Metay in STL and Pierre-André Dejaifve in ALL. MV and José Almeida (GBFor, INRAE) collected data in LAN. L. Mathieu, C. Azalbre, J. Galand and Maya Doureradjam helped with data collection in SPM in 2024. The authors acknowledge Corinne Buret and Vanina Benoit for DNA extraction and quality control for SNP genotyping and Michel Chantereau for developing the flowering scale with the collaboration of MV.

## Author contributions

Conceptualization: HD, XC Data curation: XC, JD, HD, VJ Formal analysis: XC, JD Funding acquisition: HD, MV Investigation: XC, HD, MV, VJ Writing – original draft: XC, HD Writing – review & editing: All authors.

## Data availability

Climatic, genomic and phenological data, scripts and code are available online: https://doi.org/10.57745/NEMQCH

Information on the studied genotypes is available in the GnpIS Information System via the FAIDARE data portal (https://urgi.versailles.inra.fr/faidare/), under the collection “POPULUS_NIGRA_LOIRE”.

## Supplementary material

**Supplementary figure S1.** Correlation between A) the altitudes and genetic PC2 values for each individual, B) the genetic PC1 values and the climatic PC2 for each individual.

**Supplementary figure S2.** Flowering date and end of flowering by clone across years in SPM

**Supplementary figure S3.** Ranking correlation by year and sex for beginning of flowering

**Supplementary figure S4.** Ranking correlation by year and sex for end of flowering

**Supplementary figure S5.** Ranking correlation by year and sex for flowering duration

**Supplementary figure S6.** Supplementary Figure S6 - Common flowering by year in SPM

**Supplementary figure S7.** Common flowering by population

