## Supplementary figures and images for "Studying the role of climate and sex in flowering dynamics of black poplar across time and space"

### Supplementary figure S1 - Correlation between the altitudes, genetic values and climatic PC2 for each individual

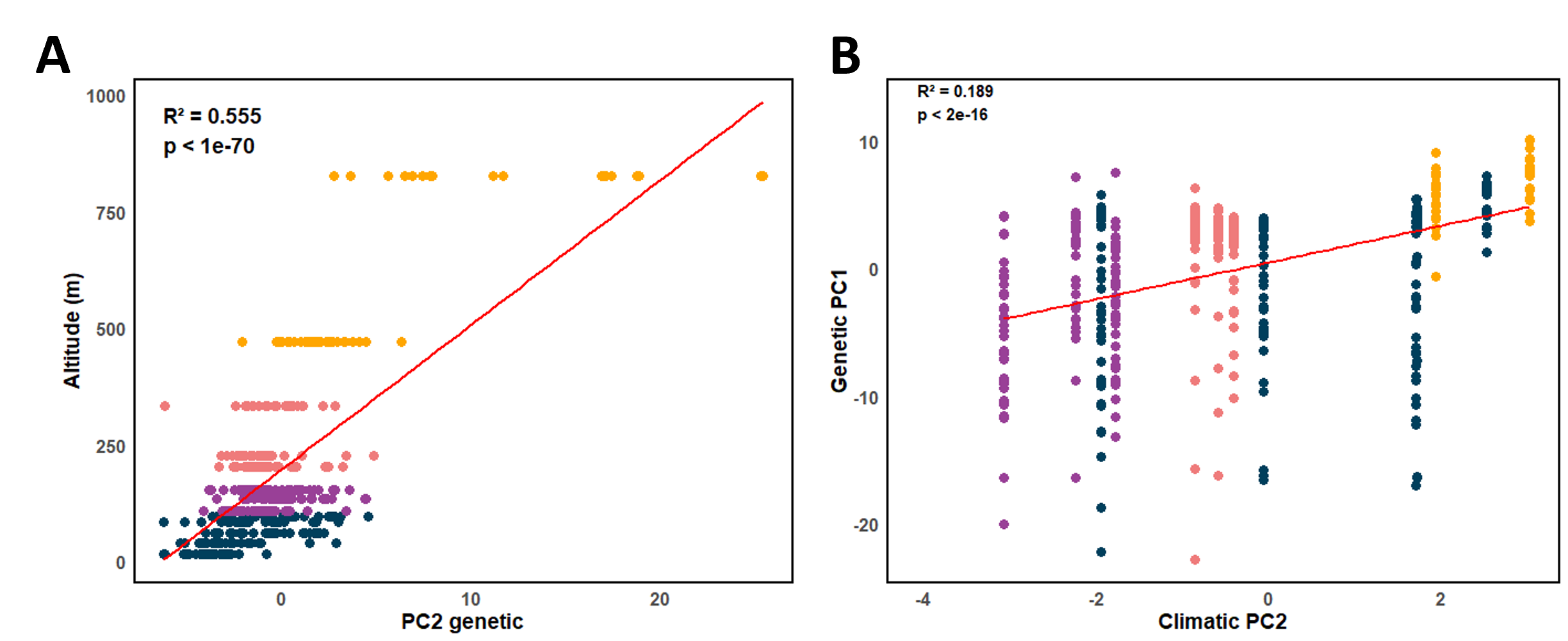
